# Interfruit : Deep Learning Network for Classifying Fruit Images

**DOI:** 10.1101/2020.02.09.941039

**Authors:** Wenzhong Liu

## Abstract

Fruit classification is conductive to improving the self-checkout and packaging systems. The convolutional neural networks automatically extract features through the direct processing of original images, which has attracted extensive attention from researchers in fruit classification. However, due to the similarity of fruit color, it is difficult to recognize at a higher accuracy. In the present study, a deep learning network, Interfruit, was built to classify various types of fruit images. A fruit dataset involving 40 categories was also constructed to train the network model and to assess its performance. According to the evaluation results, the overall accuracy of Interfruit reached 93.17% in the test set, which was superior to that of several advanced methods. According to the findings, the classification system, Interfruit, recognizes fruits with high accuracy, which has a broad application prospect.

## 1. Introduction

In the food industry, fruit represents a major component of fresh produce. Fruit sorting can not only help children and the visually impaired people to guide their diet^1^, but also can assist the or grocery stores in improving the self-checkout, fruit packaging, and transportation systems. Fruit classification has always been a relatively hard problem, which is ascribed to the wide variety and irregular shape, color and texture characteristics^2^. In most cases, the trained operators are employed to visually inspect fruits, which requires that these operators should be familiar with the unique characteristics of fruits and maintain the continuity as well as consistency of identification criteria^3^. Given the lack of a multi-class automatic classification system for fruits, researchers have begun to employ Fourier transform near infrared spectroscopy (FTIR), electronic nose, and multi spectral imaging analysis for fruit classification^4^.

The image-based fruit classification system requires only a digital camera, and achieves favorable performance, which has thus attracted considerable attention from numerous researchers. Typically, this new solution adopts wavelet entropy, genetic algorithms, neural networks, support vector machines, and other algorithms to extract the color, shape, and texture characteristics of fruits for recognition ^5^. For fruits that have quite similar shapes, color characteristics have become the criteria for the successful fruit classification^6^. Nonetheless, these traditional machine learning methods require the manual feature extraction process, and feature extraction methods may be redesigned in calibrating parameters^7^. For example, for the apple and persimmon images with extremely similar color and shape, it is difficult for the traditional methods to accurately distinguish between them. To solve this problem, a computer vision-based deep learning technology is proposed^8^. Notably, deep learning is advantageous in that, it directly learns the features of fruit images from the original data, and the users do not need to set any feature extraction method^9^. Convolutional neural networks stand for the earliest deep learning methods used for identifying fruits, in which numerous techniques, such as convolution, activation, and dropout, are adopted^10^. The existing fruit data sets are small, and insufficient training is likely to occur in the modeling process. The Weekly-Shared Deep Transfer Networks (DTN) can alleviate the deficiency of image training data^11, 12^. Researchers are working to apply the technology to other image sets. However, the deep learning methods have not been extensively utilized to classify different categories of fruits, and the classification accuracy is still not high^13^.

To enhance the recognition rate of deep learning for fruits, a deep learning architecture named Interfruit was was proposed in the present study for fruit classification, like a tree network that integrates the AlexNet, ResidualBlock, and Inception networks. Additionally, a common fruit dataset containing 40 categories was also established for model training and performance evaluation. Based on the evaluation results, Interfruit achieved superior classification accuracy to the existing fruit classification methods.

## 2. Materials and Methods

### 2.1 Data set

Totally 3,139 images of common fruits in 40 categories were collected from Google, Baidu, Taobao, and JD.com to build the image data set, IntelFruit (Figure 1). Each image was cropped to 300*x*300 pixels. Table 1 shows the category and number of fruit pictures used in the current study. For each type of fruit images, 70% images were randomly assigned to the training set, while the remaining 30% were used as the test set. The as-constructed model was trained based on the training set and evaluated using the test set.

**Table 1.** Summary of the training and test sets

| Label | Category | Number of Training Set | Number of Test set | Total Number |
| --- | --- | --- | --- | --- |
| 0 | Apple | 45 | 18 | 63 |
| 1 | Apricot | 25 | 10 | 35 |
| 2 | Avocado | 47 | 19 | 66 |
| 3 | Banana | 28 | 12 | 40 |
| 4 | Blueberry | 47 | 20 | 67 |
| 5 | Brin | 84 | 36 | 120 |
| 6 | Cantaloupe | 73 | 31 | 104 |
| 7 | Carambola | 42 | 17 | 59 |
| 8 | Cherry | 47 | 19 | 66 |
| 9 | Cherry Tomatoes | 52 | 22 | 74 |
| 10 | Citrus | 49 | 20 | 69 |
| 11 | Coconut | 94 | 40 | 134 |
| 12 | Durian | 54 | 22 | 76 |
| 13 | Ginseng fruit | 46 | 19 | 65 |
| 14 | Grapefruit | 62 | 26 | 88 |
| 15 | Grape_Black | 127 | 54 | 181 |
| 16 | Grape_Green | 41 | 17 | 58 |
| 17 | Hawthorn | 84 | 35 | 119 |
| 18 | Jujube | 98 | 41 | 139 |
| 19 | Kiwi | 31 | 12 | 43 |
| 20 | Lemon | 35 | 15 | 50 |
| 21 | Longan | 95 | 40 | 135 |
| 22 | Loquat | 51 | 21 | 72 |
| 23 | Mango | 47 | 19 | 66 |
| 24 | Mangosteen | 39 | 16 | 55 |
| 25 | Mulberry | 42 | 17 | 59 |
| 26 | Olive | 42 | 18 | 60 |
| 27 | Orange | 50 | 21 | 71 |
| 28 | Passion fruit | 65 | 27 | 92 |
| 29 | Peach | 54 | 22 | 76 |
| 30 | Pear | 26 | 10 | 36 |
| 31 | Persimmon | 45 | 19 | 64 |
| 32 | Pineapple | 115 | 49 | 164 |
| 33 | Pitaya | 82 | 35 | 117 |
| 34 | Plum | 28 | 12 | 40 |
| 35 | Prunus | 35 | 14 | 49 |
| 36 | Rambutan | 59 | 25 | 84 |
| 37 | Sakyamuni | 48 | 20 | 68 |
| 38 | Strawberry | 58 | 24 | 82 |
| 39 | Watermelon | 24 | 9 | 33 |
|  | Sum. | 2216 | 923 | 3139 |

**Figure 1.**
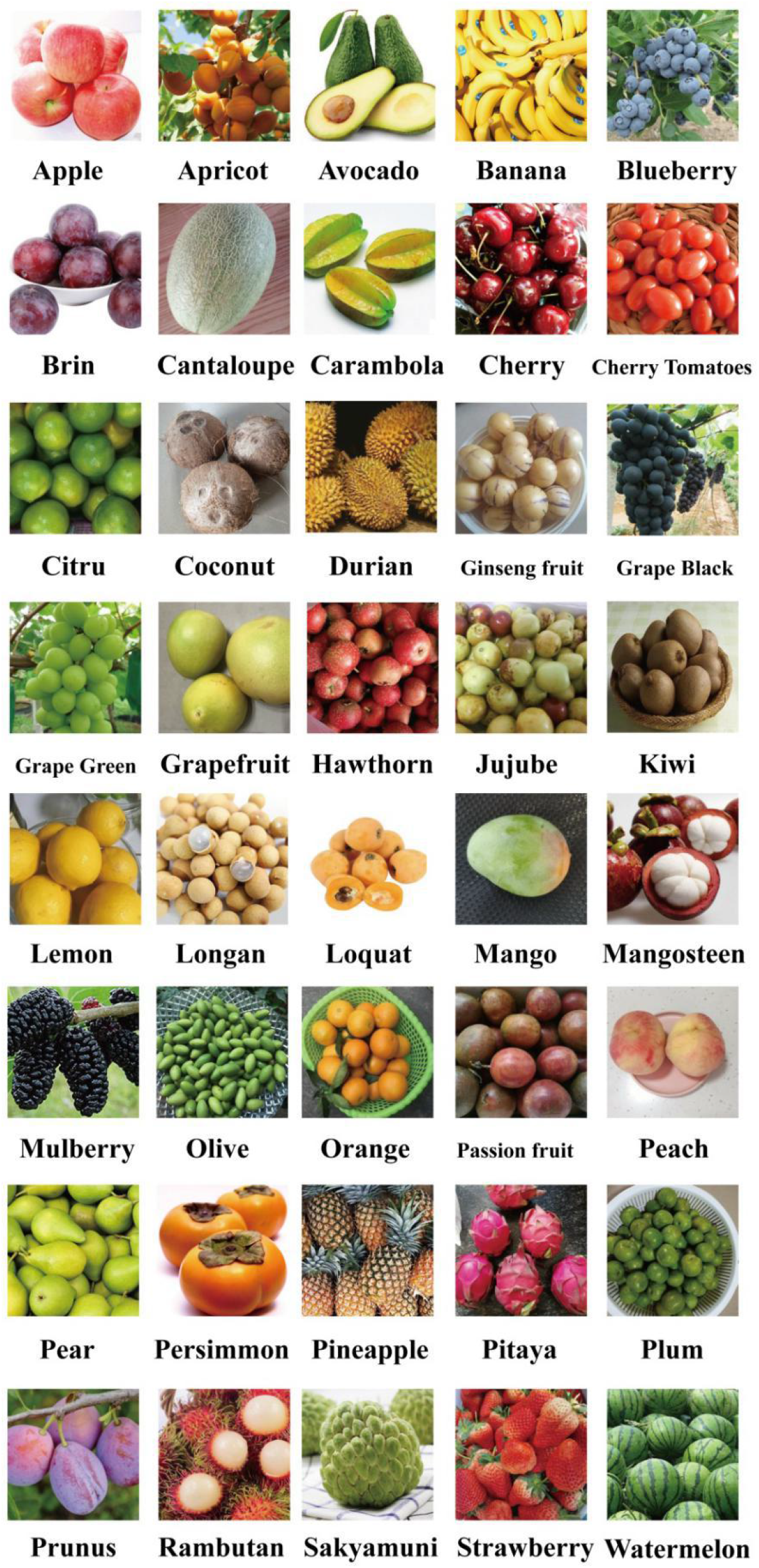
Categories of IntelFruit data set.

### 2.2 Convolutional Layer

The convolutional neural networks are a variant of deep networks, which automatically learn the simple edge shapes from raw data, and identify the complex shapes within each image through feature extraction. The convolutional neural networks include various convolutional layers similar to the human visual system. The convolutional layers generally have filters with the kernels of 11 × 11, 9 × 9, 7 × 7, 5 × 5 or 3 × 3. The filter fits weights through training and learning, while the weights extract features, just similar to the camera filters.

### 2.3 Rectified Linear Unit (ReLU)Layer

Convolutional layers are linear and can not capture the non-linear features. Therefore, a rectified linear unit (ReLU) is used as a non-linear activation function for each convolutional layer. ReLU suggests that, when the input value is less than zero, the output value will be set to zero. Using the ReLU, the convolutional layer can output the non-linear feature maps, thereby reducing the risk of overfitting.

### 2.4 Pooling Layer

The pooling layer is adopted for compressing the feature map after the convolutional layer. The pooling layer summarizes the output of the neighboring neurons, which reduces the activation map size and maintains the unchanged feature. There are two methods in the pooling layer, i.e. maximum and average pooling. In this paper, the most popular pooling strategy, namely, the maximum pooling (MP) method was adopted, which remained the maximum pooling area.

### 2.5 ResidualBlock and Inception Structure

The general convolutional neural networks like as AlexNet tend to overfit the training data and have poor performance on the actual data. Therefore, the ResidualBlock and Inception structure were used to solve this problem in this study. The Deep Residualnetwork alters several layers into a residual block (Figure 2). The residual block is composed of ReflectionPad layer, Convolutional layer (kernel 3×3, output channel 256), InstanceNorm layer, ReLU layer, ReflectionPad layer, Convolutional layer (kernel 3×3, output channel 256). The ResidualBlock Network solves the degradation problem of deep learning networks, accelerates the training speed of deep networks, and promotes the faster network convergence.

**Figure 2.**
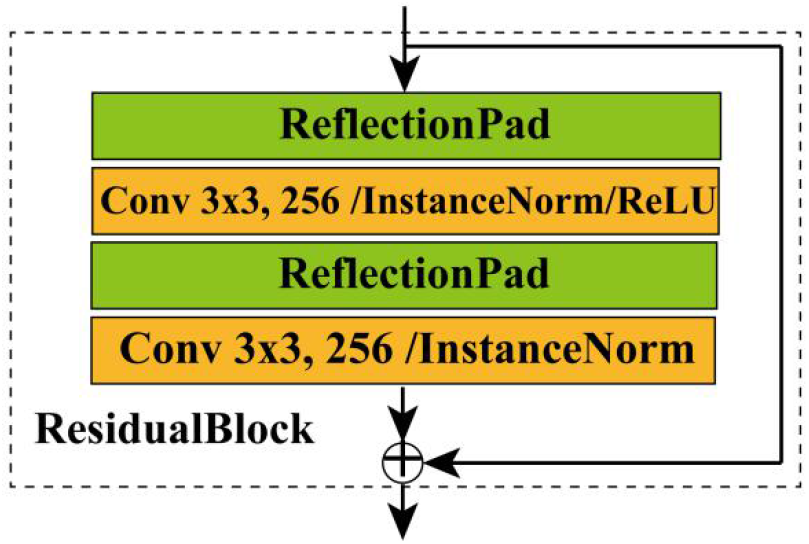
ResidualBlock Structure.

In addition, the Inception structure (Figure 3) connects the results of convolutional layers with different kernel sizes to capture the features of multiple sizes. In this study, the inception module was integrated into one layer by several parallel convolutional layers. There are four branches in the Inception structure. The first branch consists of a convolutional layer, batch normalization layer, and ReLU layer, with the kernel of the convolutional layer of 1*1 and the stride of 1. The second branch is comprised of two pairs of convolutional layers, the batch normalization layer and the ReLU layer, with the kernel of the first convolutional layer of 1*1 and the stride of 1, while the kernel of the second convolutional layer is 3*3 and the stride is 1. The third branch also consists of two pairs of convolutional layers, batch normalization layer, and ReLU layer. Of them, the kernel of the first convolutional layer is 1*1, and the stride is 1. The kernel of the second convolutional layer is 5*5, and the stride is 1. The fourth branch is made up of the pooling layer, convolution layer, batch normalization layer, and ReLU layer. The kernel of the pooling layer is 3*3 and the stride is 1. The kernel of the convolutional layer is 1*1 and the stride is 1. There are seven input parameters in the Inception. The first parameter is the input channel, and the second parameter is the output channel value of the first branch. The third parameter is the output channel value of the first layer of the second branch, and the fourth parameter is the output of the second branch. The fifth parameter is the output channel value of the first layer on the third branch and the sixth parameter is the third branch’s output channel value. The seventh parameter is the output channel value of the fourth branch. Inception reduces the size of both modules and images, and increases the number of filters. Further, the module learns more features with fewer parameters, making it easier for the 3D space learning process.

**Figure 3.**
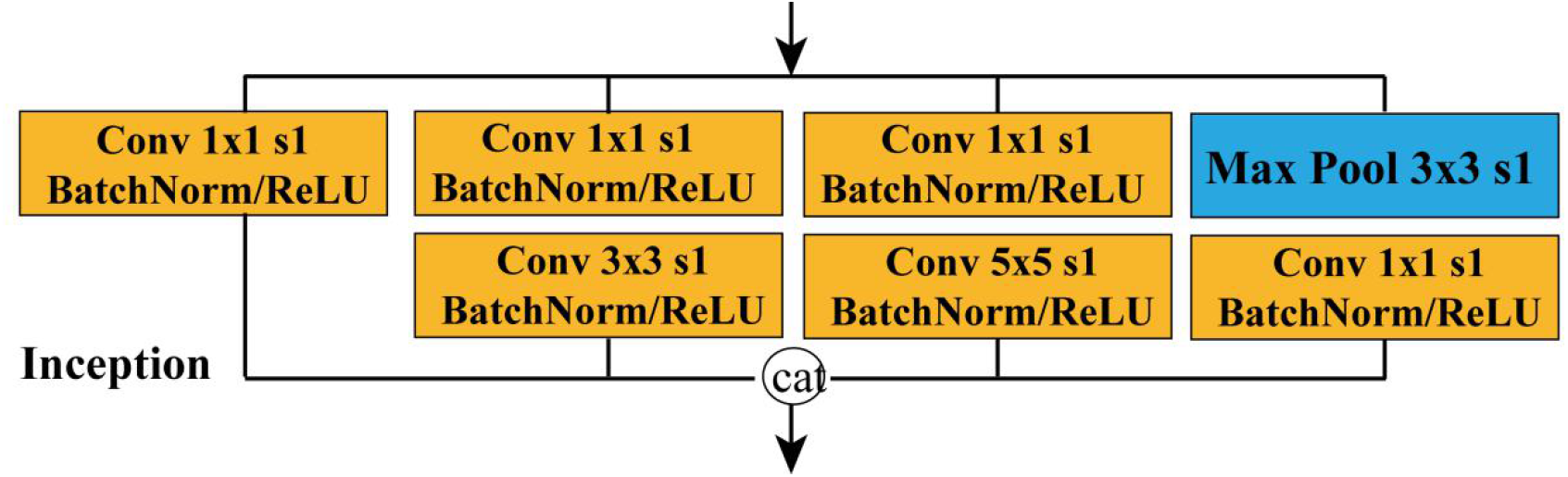
Inception Structure.

### 2.6 Fully Connected and Dropout Layer

Fully connected layer (FCL) is used for inference and classification. Similar to the traditional shallow neural network, FCL also contains many parameters to connect to all neurons in the previous layer. However, the large number of parameters in FCL may cause the problem of overfitting in the case of training, while the dropout method serves as a technique to solve this problem. Briefly, the dropout method is implemented during the training process by randomly discarding units connected to the neural network. In addition, the dropout neurons are randomly selected during the training step, and its appearance probability is 0.5. During the test step, the neural network is used without dropout operation.

### 2.7 Model Structure and Training Strategy

In this study, the convolutional neural network, IntelFruit, was constructed to classify fruits (Figure 4). According to Figure 4, the input image with a size of 227×227×3 was fed into the IntelFruit network. The IntelFruit model was a stack architecture integrating convolutional network+ ResidualBlock + Inception structure. The IntelFruit model is like an inverted trigeminal tree. The intermediate branch layers are AlexNet net, ResidualBlock structure and Inception structure. The input and output channels of the ResidualBlock structure are both 256, while those of Inception are 256 and 84, respectively. The output channels of the Inception’s branches are 16, 16, 48, 4, 12, and 8. The left branch is derived from the second convolutional layer of AlexNet. The left unit has an Inception structure and a Max Pool layer, in which the output channels of the Inception’s branches are the same as the intermediate branch. The input and output channels of Inception are 256 and 84, respectively, whereas those of Max Pool layer are 84, the kernel is 3*3, and the stride is 2. The right branch is derived from the third convolutional layer of AlexNet. The right unit has an Inception structure and a Max Pool layer with the same as the left branch. The input and output channels of Inception are 384 and 84, respectively. The output size of the three units is all 84*6*6. According to the first channel, each branch’s output is merged and then reshaped to a one-dimensional vector of 9072=84*6*6*3. The last three-level structure is the FC layer, the input length and output length of the first-level FCL are 9072 and 512, respectively; while those of the second-level FC are 512 and 512; those of the third-level FC are 512 and 40. There are ReLU and Dropout layers between FCL, and the ratio of Dropout is 0.5. Notably, the last fully connection layer played a role as a classifier, which calculated and output the scores of different fruits.

**Figure 4.**
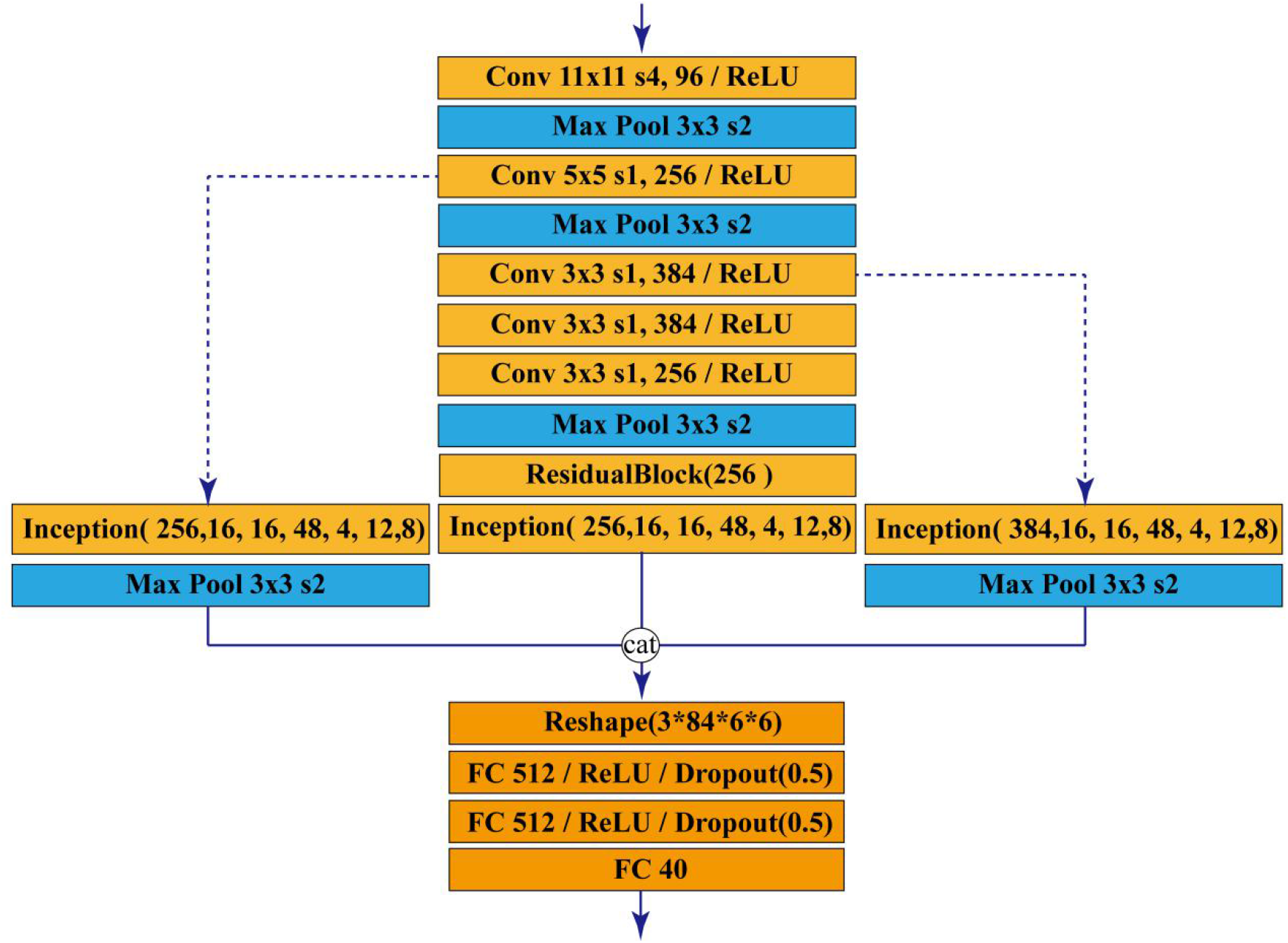
InterFruit Model Structure.

To minimize errors, the Adam optimizer was also employed in this study, which was superior in its high computational efficiency, low memory requirements and great suitability for large data or many parameters. The learning rate of the Adam optimizer was set to a constant of 1×10e-4, and CrossEntropyLoss was used as a cost function. Thereafter, the as-proposed model was trained and tested end-to-end on the CPU i5-8400processor, GPU K80 24G, with 64 GB of running memory and the operating system of WIN 10 x64.

### 2.8 Metrics of Performance Evaluation

The prediction performance of classifier was evaluated by two metrics, including accuracy (***Acc***) and average ***F1-score***. To be specific, the metrics were defined as follow:

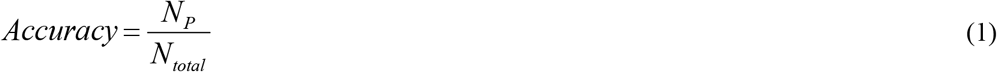

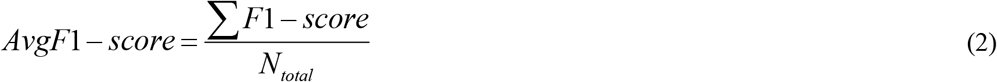

where *N*_*p*_ is the number of all correctly classified pictures and *N*_*total*_ is the number of all pictures. Average *F1-score* was calculated using the method average = “weighted”of the sklearn.metrics package

## 3. Results and Discussion

### 3.1 Loss and Accuracy Rate Curve

In terms of time and memory consumption in model training, the loss vs. accuracy curve is an effective feature. Figure 5.A presents the loss rate curve of IntelFruit on training and test sets in 200 iterations. Clearly, the loss curve of the test set was similar to the training set, with lower errors at epoch 46, indicating the high stability of IntelFruit. Figure 5.B illustrates the accuracy curves of the training and testing sets. A low higher accuracy rate was achieved at epoch 116, suggesting that IntelFruit effectively learned data and might serve as a candidate model for fruit recognition.

**Figure 5.**
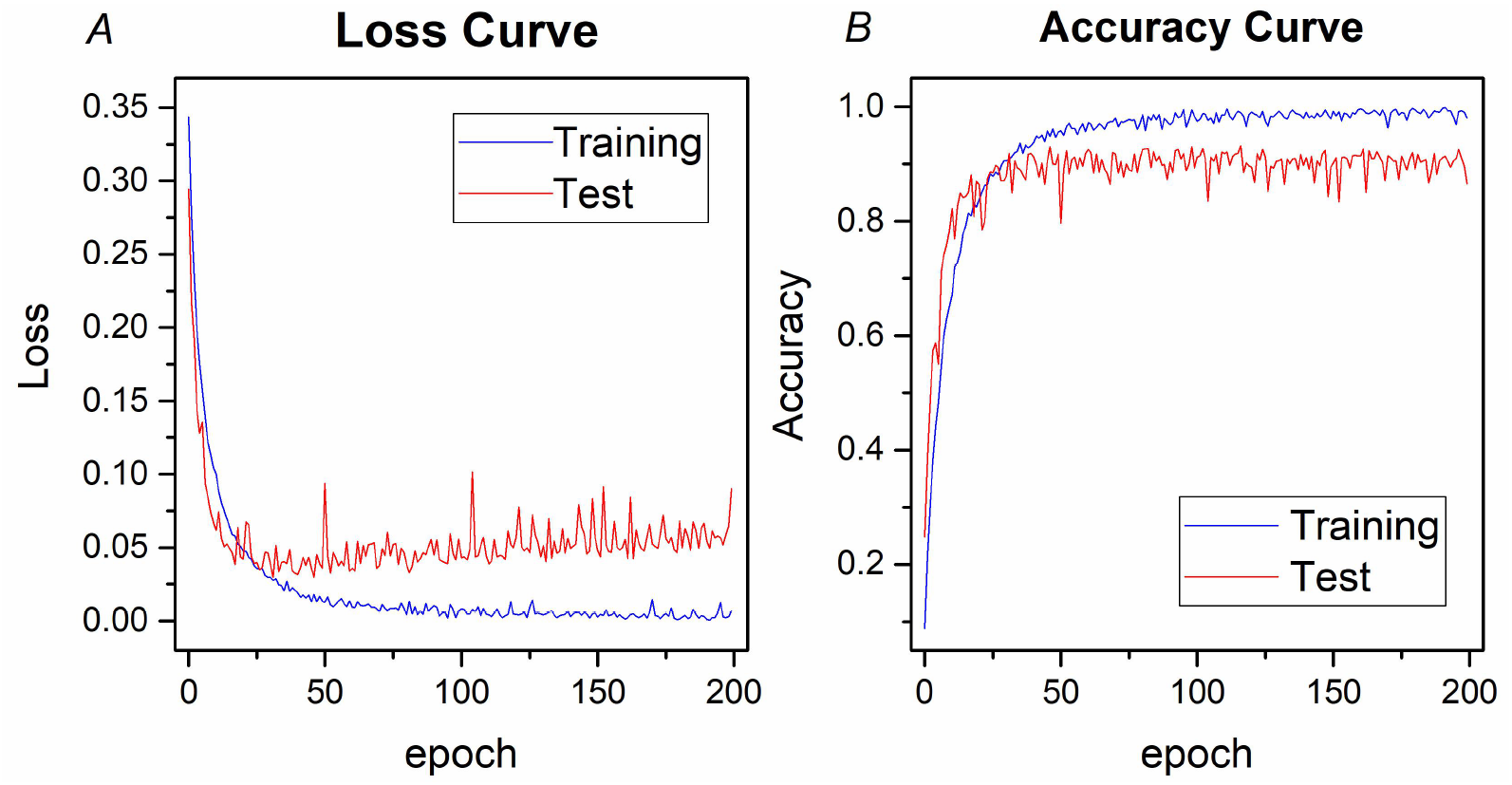
Loss and Accuracy Curves.

### 3.2 Confusion Matrix

In the current work, the proposed deep learning network IntelFruit was trained on the fruit dataset. Afterwards, the model was evaluated on the test set, showing good performance. Figure 6 displays the confusion matrix of the classification results, where each row represents the actual category while each column stands for the predicted result. In addition, the number (*m-th* row and *n-th* column) indicated the number of actual instances to the *m-th* label and was predicted as the *n-th* label.

**Figure 6.**
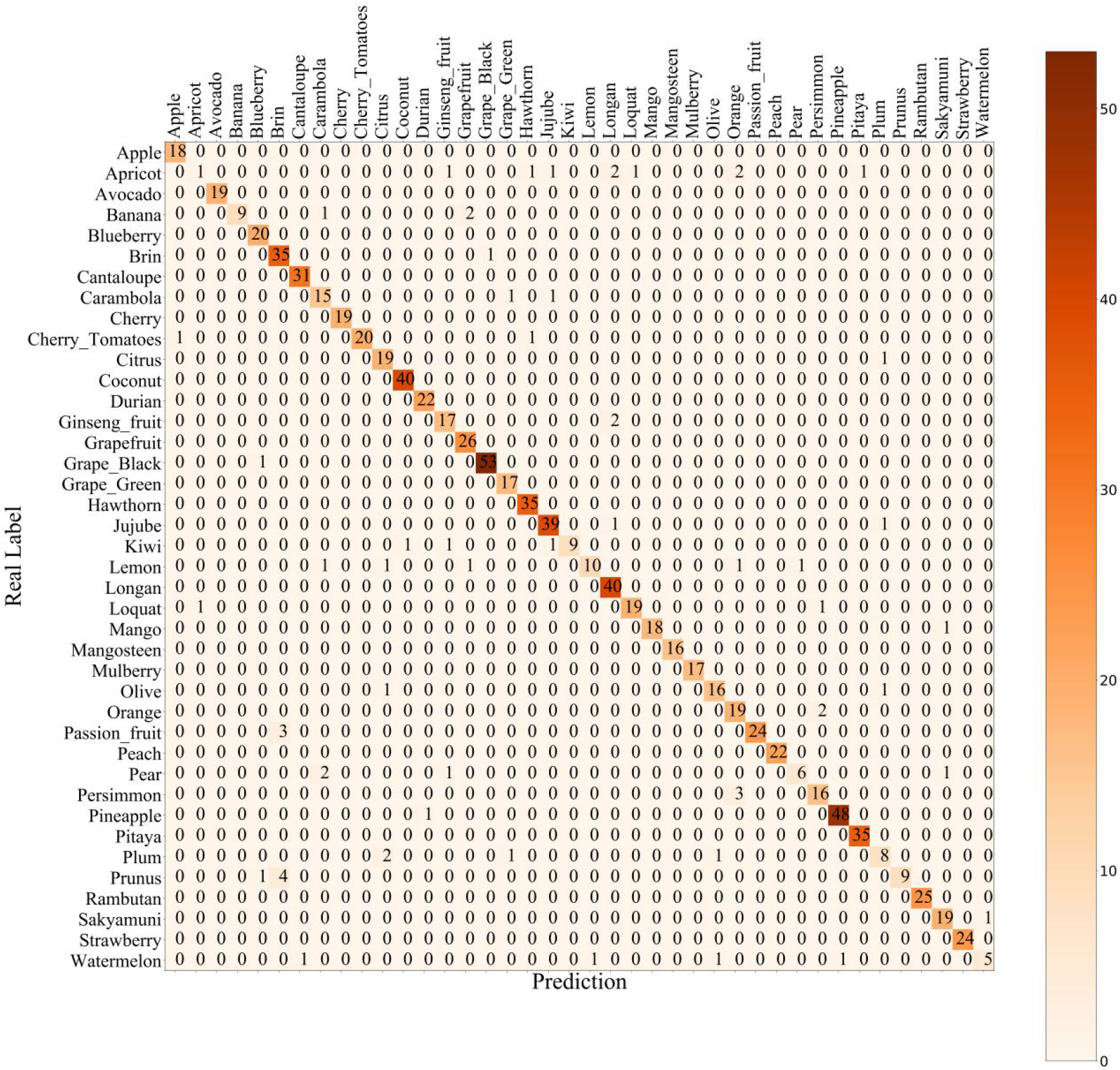
Confusion matrix on the test set.

The performance of the classifier was visually evaluated based on the results, and highlighted classes and features of the network model were also determined. IntelFruit obtained a high recognition rate. Typically, the best classified fruits were Grape_Black and Pineapple with different shapes, colors and characteristics from those of other fruits. As clearly observed from Figure 6, among the 40 categories, 63 images were incorrectly predicted as the 27 category, and the remaining 860 images were correctly predicted. Therefore, the best classified fruits were Grape_Black and Pineapple, closed to a high accuracy of 100%. By contrast, the worst classified fruits were Apricot and Plum, with a low accuracy. According to the above results, the IntelFruit model was able to better identify different fruits.

### 3.3 Comparison of Classification Performance

To evaluate the effectiveness of these models, the as-proposed method was compared with the existing methods for modern deep learning. The models were evaluated on the test set by the accuracy rate, and avg *F1-score* (Table 2). In Table 2, the model IntelFruit achieved lower false positive and false negative rates, which demonstrated the effectiveness. For the fruit dataset involving 40 categories, the accuracy value of the proposed model was 93.17%, which was subsequently compared with those of AlexNet, GoogLeNet and ResidualBlock18. The accuracy values of these three methods were as follows, 83.97% for AlexNet, 84.83% for GoogLeNet, and 75.52% for ResidualBlock18. In addition, Table 2 clearly illustrated that the avg F1-score of the proposed model was 96.47%, which was also superior to the existing models.

**Table 2.** Comparison of Classification Performance by training on Intelfruit dataset

| Methods | *Acc* | *F1score* |
| --- | --- | --- |
| AlexNet | 83.97 | 91.28 |
| GoogLeNet | 84.83 | 91.79 |
| ResidualBlock18 | 75.52 | 86.05 |
| Intelfruit | 93.17 | 96.47 |

It also allowed for the transferred learning using the AlexNet, GoogLeNet, and ResidualBlock18 models. The trained AlexNet, GoogLeNet, and ResidualBlock18 models were downloaded from the official website of pytorch. The last FC layer’s output was changed to 40 categories, and the default weights were set the other layers. Then, the data set in this paper was used for training and prediction. Table 3 shows the prediction results. The accuracy values of these three methods were as follows, 83.21% for AlexNet, 83.97% for GoogLeNet, and 76.60% for ResidualBlock18. Besides, avg F1-score values of these three methods were 90.83% for AlexNet, 91.28% for GoogLeNet, and 86.74% for ResidualBlock18. The accuracy and F1-score of these three methods were lower than those of IntelFruit too. It was seen from Table 2 and Table 3 that, the transfer learning method had not greatly improved the prediction metrics of these three methods.

**Table 3.** Comparison of Classification Performance by transfer learning

| Methods | *Acc* | *F1score* |
| --- | --- | --- |
| AlexNet | 83.21 | 90.83 |
| GoogLeNet | 83.97 | 91.28 |
| ResidualBlock18 | 76.60 | 86.74 |
| Intelfruit | 93.17 | 96.47 |

In the case of fruit recognition, IntelFriu was more effective than those previous methods, revealing the superiority of the proposed network. In general, the intelFriut model with the highest recognition rate shows promising application value in the food and supermarket industries. Noticeably, IntelFruit was associated with many advantages, and it ushered in a new method to classify 40 different types of fruits simultaneously. According to the high-precision results, convolutional neural networks might also be used to achieve high performance and faster convergence, even for the smaller data sets. This model captured images to train the model without preprocessing the images to eliminate the background noise and the lighting settings. Although this model showed excellent performance in the evaluated cases, it was linked with some difficulties in some cases. For instance, for categories Apricot and Plum, some categories were easily confused with others due to the insufficient sample sizes, leading to false positives or lower accuracy.

## 4. Conclusions

It is quite difficult for the supermarket staff to remember all the fruit codes, and it is even more difficult to sort the fruits automatically if no barcodes are printed on the fruits. In the current work, a novel deep convolutional neural network named intelFriut is proposed, which is then used to classify the common fruits and help supermarket staff to quickly retrieve the fruit identification ID and price information. IntelFriut is an improved stack model that integrates AlexNet, ResidualBlock and Inceptiont, with no need for extracting color and texture features. Furthermore, different network parameters and DA techniques are used to improve the prediction performance of this network. Besides, this model is evaluated on the fruit dataset intelFriut, and compared with several existing models. The evaluation results demonstrate that the intelFriut network proposed in the present study achieves the highest recognition rate, with the overall accuracy of 93.17%, which is superior to other models. Findings in the present study indicate that the network combining AlexNet, ResidualBlock and Inception achieves higher performance and has technical validity. Therefore, it can be concluded that, intelFriut is a novel and highly computational tool for fruit classification with broad application prospects.

## Availability of data and materials

All data sets and codes used in this study are available at https://github.com/lwzyb/Interfruit.

